# PREDICTING SPEECH INTELLIGIBILITY FROM A SELECTIVE ATTENTION DECODING PARADIGM IN COCHLEAR IMPLANT USERS

**DOI:** 10.1101/2021.09.17.460821

**Authors:** Waldo Nogueira, Hanna Dolhopiatenko

**Affiliations:** Medical University Hannover, Cluster of Excellence “Hearing4all”, Hannover, Germany

## Abstract

1.

**Objectives:** Electroencephalography (EEG) can be used to decode selective attention in cochlear implant (CI) users. This work investigates if selective attention to an attended speech source in the presence of a concurrent speech source can predict speech understanding in CI users.

**Approach:** CI users were instructed to attend to one out of two speech streams while EEG was recorded. Both speech streams were presented to the same ear and at different signal to interference ratios (SIRs). Speech envelope reconstruction of the to-be-attended speech from EEG was obtained by training decoders using regularized least squares. The correlation coefficient between the reconstructed and the attended (*ρ_A_SIR__*) and between the reconstructed and the unattended (*ρ_U_SIR__*) speech stream at each SIR was computed.

**Main Results:** Selective attention decoding in CI users is possible even if both speech streams are presented monaurally. A significant effect of SIR on the correlation coefficient to the attended signal *ρ_A_SIR__*, as well as on the difference correlation coefficients *ρ_A_SIR__* – *ρ_U_SIR__* and *ρ_A_SIR__* – *ρ_U_SIR__* was observed, but not on the unattended correlation coefficient *ρ_U_SIR__*. Finally, the results show a significant correlation between speech understanding performance and the correlation coefficients *ρ_A_SIR__–ρ_U_SIR__* or −*ρ_U_SIR__* across subjects. Moreover, the difference correlation coefficient *ρ_A_SIR__* – *ρ_U_−SIR__*, which is less affected by the CI electrical artifact, presented a correlation trend with speech understanding performance.

**Significance:** Selective attention decoding in CI users is possible, however care needs to be taken with the CI artifact and the speech material used to train the decoders. Even if only a small correlation trend between selective attention decoding and speech understanding was observed, these results are important for future development of objective speech understanding measures for CI users.

## 2. INTRODUCTION

A cochlear implant (CI) is a small electronic medical device that can restore hearing to people suffering from profound hearing loss. The CI bypasses the damaged structures of the inner ear by directly stimulating the auditory nerve. Although most CI users obtain good speech intelligibility in quiet environments, their speech recognition deteriorates in environments containing noise or competing talkers (e.g. Cullington and Zeng, 2008). For this reason, it is essential to program the CI to optimize its outcome. The outcome is usually measured as the number of correct understood words from a clinical speech test, in which words are presented and the CI user is asked to repeat the recognized words. However, these tests are difficult to perform in people with fluctuating engagement (i.e. children) or with absent response (i.e. unconscious patients or subjects with additional disabilities). Moreover, such behavioral tests are provided by an audiologist, making them time consuming and subjective because these depend on the examiner’s experience. For this reason, several objective paradigms have been proposed to predict the speech performance of CI users that do not depend on the experience of the audiologist and that do not require the behavioral response of the tested subject (e.g. Gransier et al., 2016; Verschueren et al., 2018). Such paradigms could find application to speed up the fitting procedure, to automatically optimize the speech performance of CI users and to track the speech performance of CI users objectively across time.

In this context, several objective peripheral and cortical measures based on electroencephalography (EEG) have been proposed to assess hearing performance. These measures include auditory brainstem responses (ABRs; e.g. Verhaert et al., 2008; Anderson et al., 2013), auditory steady state responses (ASSRs; e.g. Picton et al., 2005; Luts et al., 2006; Gransier et al., 2016), cortical auditory evoked potentials (CAEPs: e.g. Sharma and Dorman, 2006; Liebscher et al., 2018), acoustic change complex (ACC; e.g. Undurraga et al., 2021) or P300/mismatch negativity (MMN; e.g. Martin et al., 2008; Roman et al., 2005). More specifically, ASSRs have been found to highly correlate with speech understanding performance in CI users (Gransier et al., 2020), however its predictive power may be limited by the use of simple auditory stimuli (Lesenfants et al., 2019).

For this reason, it has been suggested that measures of brain activity based on EEG neural tracking to natural running speech can provide a more realistic and objective measure of speech understanding performance (e.g. Ding and Simon, 2012; Vanthornhout et al., 2018). In neural tracking, the brain responses can be decoded from the speech signal with the so called forward model (Haufe et al., 2014). Using this method, a temporal response function (TRF) can be derived in which interpretable peaks, such as the N1 and the P2 can be observed (Lalor et al., 2009; Petersen et al., 2016; Di Liberto et al., 2018). In contrast, the backward model can be used to reconstruct the speech envelope from the recorded EEG. The correlation coefficient between the original and the reconstructed speech stream is used as a measure of neural coding of speech. Previous studies used neural tracking methods in which the listener was asked to attend and to understand the speech uttered by a single speaker. Recently, it was proposed that the correlation coefficient derived from a neural tracking paradigm can predict speech understanding performance in normal hearing (NH) listeners (e.g. Ding and Simon, 2013; Ding et al., 2014; Di Liberto et al., 2018; Vanthornhaut et al., 2018, Lesenfants et al., 2019, Vanthornhaut et al., 2019, Decruy et al., 2020). Indeed, it was shown that higher correlation coefficients are observed when the presented speech is more understandable for the listeners (Etard et al., 2019; Dimitrijevic et al., 2019). More recently, it has been shown that the correlation coefficient of neural tracking can predict speech understanding performance in CI users (Paul et al., 2020). However, it is necessary to suppress the CI artifact (Sommers et al., 2019). The CI stimulates the auditory nerve following a stimulation pattern that resembles the speech envelope. The speech envelope is therefore transmitted through slow temporal modulations that leak into the EEG that can obscure the neural recordings (e.g. Somers et al., 2018; Deprez et al., 2017; Hoffman and Wouters, 2010). Even if some methods have been proposed to reduce the CI artifact for neural tracking paradigms to single speakers, the artifact cannot be completely removed (Sommers et al., 2019).

Neural tracking can also be measured in a competing talker situation using a so called selective attention paradigm. In this case, the listener is asked to attend to one speaker and ignore all others. Previous studies have shown the possibility to decode selective attention using the forward or the backward model in NH listeners (O’Sullivan et al., 2012, Mirkovic et al, 2015) and in CI users (Nogueira et al., 2019a; Nogueira et al. 2019b; Paul et al., 2020). A potential advantage of the selective attention paradigm is that it offers the possibility to analyze relative changes in neural activity between the attended and the unattended speech stream. These relative changes in neural activity can be computed as the difference between the attended and the unattended correlation coefficients. If both speech streams are properly balanced, it can be assumed that the contribution of each speech stream to the CI artifact will be similar and that the difference between the correlation coefficients will cancel out its effects to some extent. In addition, it can also be hypothesized that the difference in correlation coefficients may be less influenced by the quality of the recorded EEG, which depends, among other factors, on the electrode impedances. Based on these hypotheses, this work will investigate if the attended correlation coefficient as well as the difference between the attended and the unattended correlation coefficients derived from a selective attention paradigm can predict speech understanding in NH listeners and CI users. In this regard, it has been shown that maximum differences between the attended and the unattended correlation coefficients occur at latencies ranging from 200 to 400 ms after stimulus onset for CI users (Nogueira et al., 2019a; Nogueira et al., 2019b). At these later lags the effect of the artifact is smaller than at earlier lags potentially enabling the use of a selective attention paradigm to predict speech understanding. Moreover, Paul et al. (2020) investigated whether neural differentiation of attended and ignored speech at an early cortical processing stage of 150 ms or at a late cortical stage of 250 ms can predict speech intelligibility in a group of NH people and bilateral CI users presenting a digit-in-noise test dichotically. Enhancement of the attended speech was observed near ~150 ms for NH controls, whereas the representation of ignored speech was stronger in bilateral CI users near ~250 ms. Furthermore, Paul et al. (2020) showed that speech understanding performance can be significantly predicted in bilateral CI users, but not in NH individuals because the behavioural task was too easy for them causing ceiling effects. So far, to our knowledge, selective attention decoding in CI users has only been investigated in bilateral CI users presenting concurrent speech streams dichotically. However, CI outcome measures in the clinic consist of monaurally presented speech. It remains a question whether it is possible to decode the attended speaker when both speech streams are presented to the same CI side. Moreover, these previous studies with CI users did not investigate the effect of changing the relative presentation levels between the attended and the unattended speech streams (signal-to-interference ratio; SIR) on selective attention decoding. Finally, these previous studies used different speech materials to train and test the selective attention decoders or used speech materials that are not typically administered in the clinical routine.

In this work we investigate whether it is possible to predict speech intelligibility of monaurally presented speech in the presence of a concurrent interfering speech stream in CI users at different SIRs. For the estimation of the selective attention correlation coefficient, the backward model was used. Besides this, the impact of the training material and the gender of the attended talker on selective attention decoding was investigated.

## 3. MATERIAL AND METHODS

### 3.1. PARTICIPANTS

10 CI users participated in the study. All patients were native German speakers. CI users were recruited from the clinical database of the Hannover Medical School (MHH) based on their clinical speech intelligibility scores in quiet and in noise. All CI users had more than 6 months experience with their device at the time of measurement. Average age for CI users was 47.4 years (ranging from 24 to 75). More details about the CI users that participated in the study are provided in Table 1. Prior to the experiment, all participants provided written informed consent and the study was carried out in accordance with the Declaration of Helsinki principles, approved by the Ethics Committee of the Hannover Medical School.

**Table 1:** Subject data with ID, gender, age at testing for the present study, implant, sound processor, duration of device use, etiology of deafness and clinical speech-in quiet scores using the Hochmair Schulz Moser (HSM) sentence test. M=male, F=female, AB=Advanced Bionics,.

| ID | Gender | Age at testing (years) | Implant | Sound processor | Duration of device use (years) | Etiology of deafness | HSM speech-in-quiet scores |
| --- | --- | --- | --- | --- | --- | --- | --- |
| CI 1 | M | 65 | AB | Harmony | 12 | Unknown | 66.03% |
| CI 2 | M | 24 | AB | Naida CI Q90 | 3 | Unknown | 85.84% |
| CI 3 | M | 35 | AB | Naida CI Q90 | 13 | Unknown | 100% |
| CI 4 | M | 52 | AB | Naida CI Q90 | 19 | Genetic | 97.16% |
| CI 5 | M | 28 | Cochlear | CP910 | 26 | Meningitis | 100% |
| CI 6 | F | 49 | AB | Naida CI Q90 | 10 | Unknown | 89.62% |
| CI 7 | M | 59 | Cochlear | CP1000 | 9 | Unknown | 99.05% |
| CI 8 | M | 28 | Cochlear | CP910 | 26 | Unknown | 94.33% |
| CI 9 | M | 75 | AB | Naida CI Q90 | 13 | Genetic | 71.7% |
| CI 10 | M | 59 | MEDEL | Opus2 | 7 | Sudden Hearing Los | 93.9% |

### 3.2. BEHAVIORAL SPEECH UNDERSTANDING

The first experiment consisted of a speech intelligibility test. The speech material was based on the original German Hochmair Schulz Moser (HSM) sentence test (Hochmair-Desoyer et al., 1997) uttered by a male voice and a recorded version of the HSM test uttered by a female voice.

Each HSM list contains 20 sentences. Each participant was asked to recall the HSM sentences from a target speaker in the presence of a competing voice. When the target talker was a male voice the competing talker was a female voice and vice versa. After each sentence a short silent pause was inserted to give enough time for the participants to spell out the sentence. As a measure of speech understanding, the percentage of correct repeated words for each condition was calculated.

In total, 20 lists were presented, 10 with the male target voice and the other 10 with the female voice. Each four lists (2 with male target and 2 with female target) were presented at a given SIR. In total, five different SIRs were used (SIR=+20dB, +15dB, +10dB, +5dB, 0dB). The sentences at negative SIRs were not provided to CI users, as those SIRs would have been too challenging for CI users and the speech score would have been 0. The SIR was manipulated by modifying the level of the interference while keeping the level of the target speaker fixed. The speech material was administered only to one ear. The best hearing side of the each subject was selected for the audio presentation. The sound was provided via direct input cable between the audio system and the speech processor for the CI users. Stimulus presentation was controlled by the Presentation Software (Neurobehavioral Systems, Inc., Berkeley, CA, United States; version 20.1) as in our previous work (Nogueira et al., 2019). Every participant adjusted the presentation level of the speech material to obtain a moderate loudness perception by means of a ten-point loudness-rating scale (where 1 was “very soft”, 7 was equivalent to “comfortably loud” and 10 was “extremely loud”).

### 3.3. DECODING SELECTIVE ATTENTION FROM SINGLE TRIAL EEG

#### 3.3.1. EEG PROCEDURE

The second experiment consisted of continuous EEG recorded while speech stimuli were presented to the study participants. Each participant was instructed to sit relaxed and calm, keep their eyes open, and maintain their gaze fixed to the front by looking to a cross displayed on a screen placed in front of the subject. High density EEG data was recorded in an electromagnetically and acoustically shielded booth using a BrainAmp System (Brain Products GmbH, Gilching, Germany) with 96 Ag/AgCl electrodes mounted in a customized, infracerebral electrode cap with an equidistant electrode layout (Easycap GmbH, Herrsching, Germany). The nose tip of the study participant served as reference while the ground electrode was placed on a central frontopolar site. Data were recorded with a sampling rate of 500 Hz and an online filter from 0.2 to 250 Hz. Impedances were controlled and maintained below 10 kΩ. A delay of 77 ms between the onset of the stimulus and the trigger captured by the BrainAmp EEG system was observed and corrected for the analysis.

The stimuli consisted of lists of sentences from the HSM test, different to the ones used during the speech intelligibility test, but with additional SIR conditions. In total, 9 SIRs were presented: −20dB, −15dB, −10dB, −5dB, 0dB, +5dB, +10dB, +15dB, +20dB. Each SIR included 4 lists (2 lists with the male voice and 2 lists with the female voice as target) except for the 0 dB SIR which included 8 lists resulting in a total of 40 lists. For positive SIRs, the SIR was manipulated by modifying the level of the interference while keeping the level of the target speaker fixed. In contrast, for negative SIRs, the SIRs were obtained by modifying the level of the target speaker while keeping fixed the level of the interferer. The EEG procedure was conducted without requiring repetition of the words by the study participants. Moreover, EEG to a 16 minutes long story (‘A drama in the air’ by Jules Verne) narrated by a single male voice was recorded. This additional recording was used to increase the amount of data to train the decoder. The speech material was presented monaurally to the better hearing side of the CI users. The CI sound processor was moved from their usual position (behind the ear) to the participant’s collar. The change in placement did not affect the participant’s sound perception with the CI, as the sound was delivered via a direct audio cable.

#### 3.3.2. EEG ARTIFACT REJECTION

Artifact rejection based on the independent component analysis (ICA) was applied to the EEG collected from CI users. The analysis was provided within the EEGLAB Toolbox (version 19.0.0b) according to the two-steps procedure described by Viola et al. (2012). As a first step Infomax ICA weights were calculated and components that presented a physiological artifact were manually determined and rejected. The second step included the determination and rejection of the electrical artifact caused by the CI. Afterwards, the data was epoched in consecutive intervals corresponding to the length of each list, resulting in 40 trials for each subject.

#### 3.3.3. EEG AND SPEECH ENVELOPE PREPROCESSING

The EEG was preprocessed in MATLAB 8.1.0.604 (R2018b; Mathworks, Natick, MA, United States) by filtering it using a bandpass filter from 2 to 8 Hz and down sampling to 64 Hz following the same procedure as Nogueira et al. (2019). The envelopes of the original audio material (HSM and story book) were estimated based on the Hilbert transform of the original audio signal. The audio signals likewise were low-pass filtered (8 Hz) and down sampled to 64 Hz.

Speech reconstruction from EEG was obtained through the backward model of neural responses (mTRF; Crosse et al., 2016). The linear modeling performs a least square estimation between neural data and the stimulus to reconstruct an envelope of the signal. The reconstructed envelope was correlated to the envelope of the original attended audio stream (attended correlation coefficient; *ρ_A_*) and to the original unattended audio stream (unattended correlation coefficient; *ρ_U_*). The correlation coefficients were calculated for different lags, representing the time shifted versions of the EEG signal. The lag window was set to 16 ms and spanned the interval from 16 ms to 520 ms.

Different decoders were designed to investigate the impact of the training material. In total three decoders were designed based on the speech material: HSM (Decoder_HSM_), the story book (Decoder_Story_) and the combination of both (Decoder_HSM+Story_). In order to investigate the impact of the uttered voice on the correlation coefficients, two additional decoders based only on male target sentences or only on female target sentences were created. The Decoder_HSM_ and the Decoder_HSM+Story_ were tested using leave-one-out cross-validation in which 1 HSM list was used for prediction and all others for training. The Decoder_Story_ was tested using the same HSM lists as the other two decoders.

### 3.4. STATISTICAL ANALYSIS

To investigate the effect of the training material, the effect of gender and the effect of SIR on the attended correlation coefficients, the difference between attended and unattended correlation coefficients for the same (*ρ_A_SIR__-ρ_U_SIR__*) and opposite SIRs (*ρ_A_SIR__-ρ_U_−SIR__*), a repeated measures ANOVA analysis as implemented in the SPSS Software version 27 (IBM, Armonk, New York, USA) was conducted. As a post-hoc test, the Wilcoxon signed-rank test was applied.

## 4. RESULTS

### 4.1. SPEECH UDNERSTANDIG PERFORMANCE

Figure 1 presents the behaviorally measured speech understanding scores for the CI users that participated in the study.

**Figure 1:**
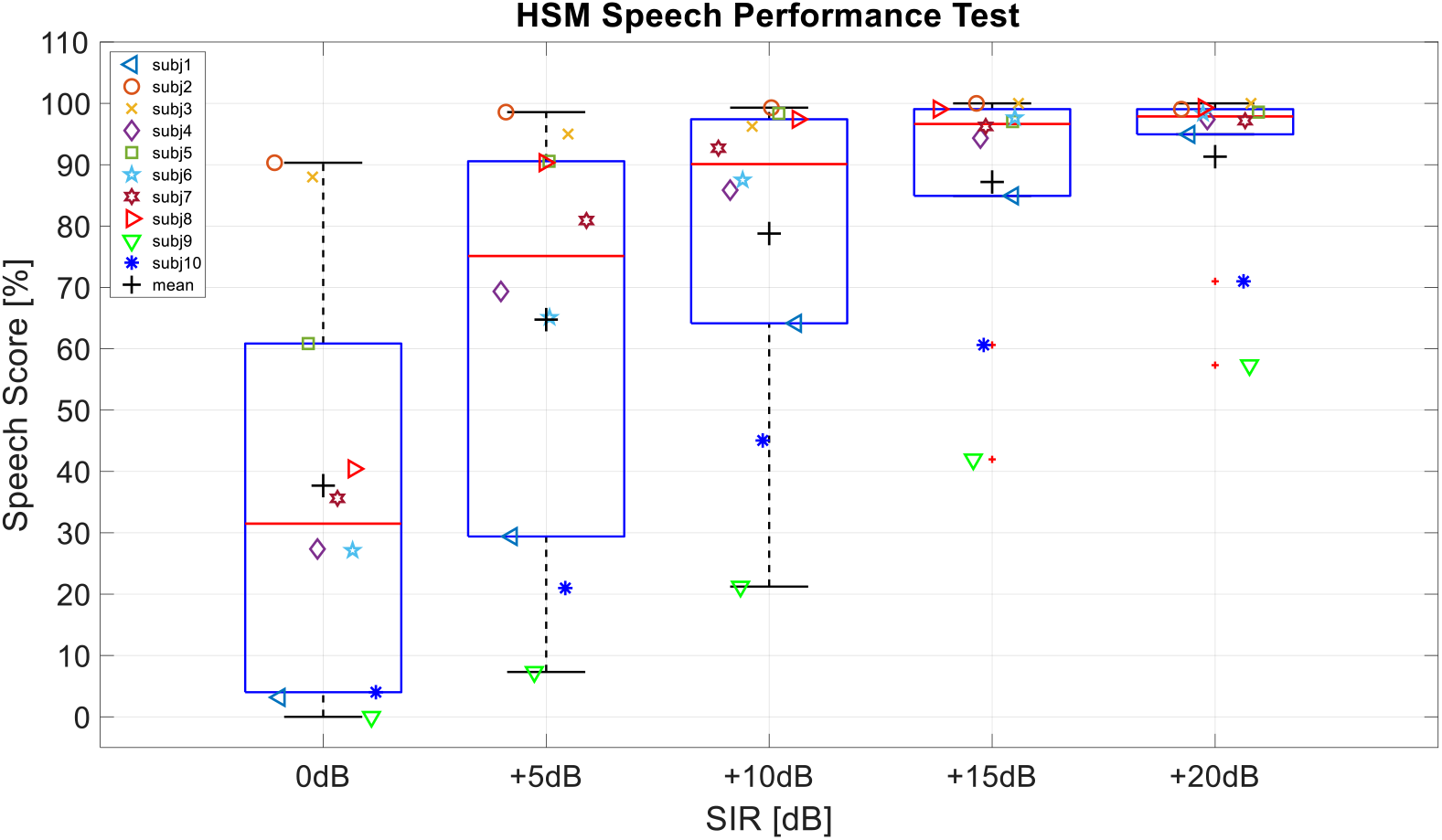
Speech understanding scores in percentage [%] correct words for each cochlear implant (CI) user and for different signal to interference ratios (SIRs) ranging from 0 dB to 20 dB. The red horizontal line in the box plots indicates the median value and the black cross indicates the mean value across subjects. The bottom and top of the box marks represent the 25^th^ and 75^th^ percentile of the speech scores across CI users.

A one way repeated measures of ANOVA revealed a significant effect of SIR on the speech understanding scores (F(4,36) = 28.988; p<0.0001). A post-hoc Wilcoxon signed rank test showed that all pairs were significantly different from each other after Bonferroni correction, except for the pair (SIR=+20 dB and SIR = +15 dB).

### 4.2. SELECTIVE ATTENTION DECODING IN COCHLEAR IMPLANT USERS

#### 4.2.1. EFFECT OF TRAINING MATERIAL ON SELECTIVE ATTENTION DECODING

Three different decoders were created using different training materials. The first one was based on the HSM material (Decoder_HSM_), the second one was based on a story book (Decoder_Story_) and the third one was based on the combination of both (Decoder_HSM+Story_). The three decoders were used to investigate the impact of training material on the attended and unattended correlation coefficients. Figure 2 presents the attended and unattended correlation coefficients across lags (Δ) for each decoder and for different SIRs after applying ICA for CI artifact reduction^1^. The ICA rejection caused reduction of the unattended correlation coefficients, especially at negative SIRs. For comparison, data in Appendix 2 presents the correlation coefficients across lags in NH listeners. As expected, the attended correlation coefficients are lower for NH listeners than for CI users, probably because the CI transmits envelope information in each electrode and the CI stimulation leaks into the EEG as artifact.

**Figure 2:**
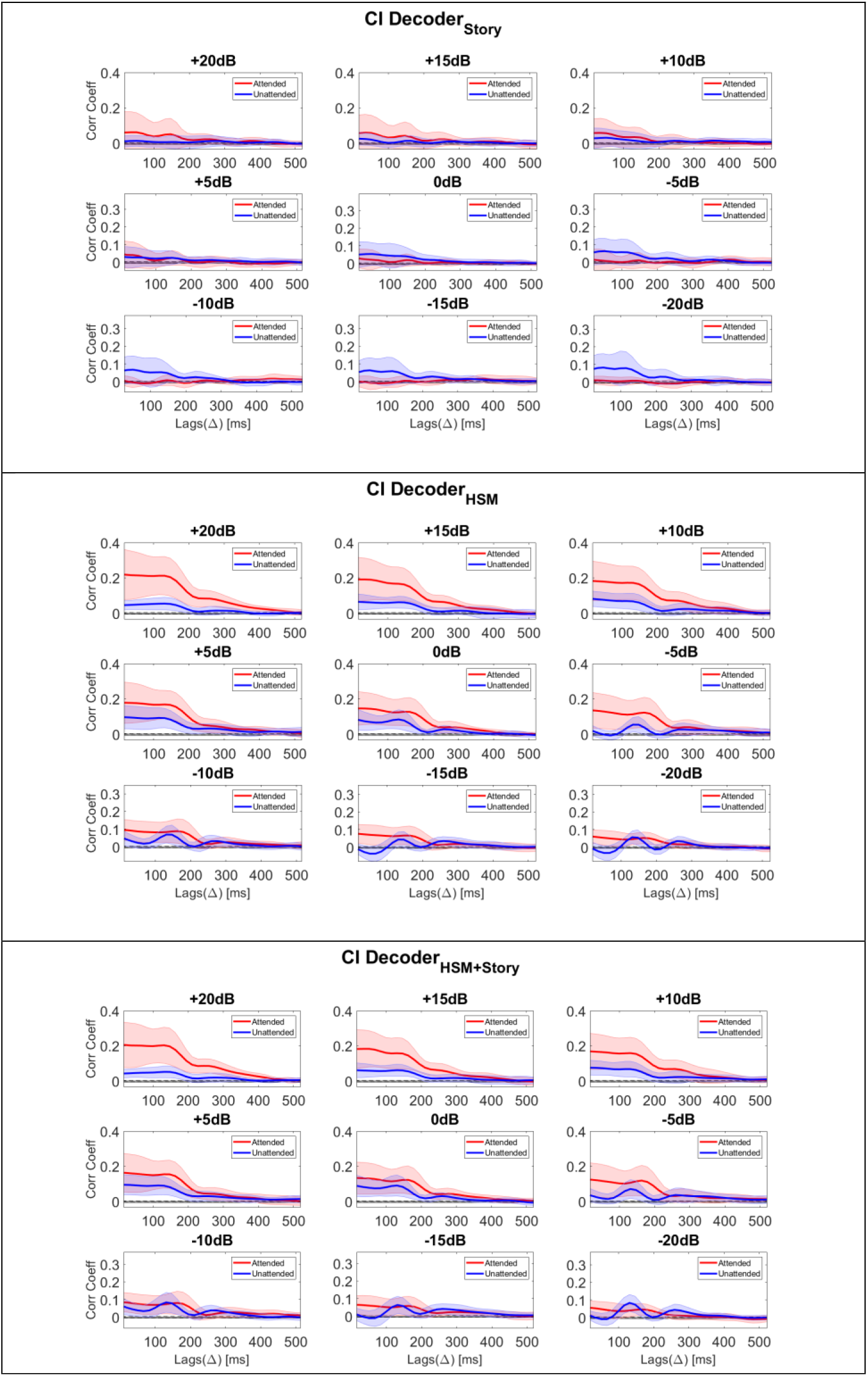
Correlation coefficients (Corr Coeff) across lags (Δ) to the attended and the unattended speech stream for cochlear implant (CI) users at different signal to interference rations (SIRs) and for different training materials: (top) Story book, (center) HSM test and (bottom) Story book + HSM test. Selective attention was decoded for SIRs ranging from −20 dB to +20 dB SIR.

In Figure 2, the morphology of the curve representing the attended correlation coefficient presents large values above 0.1 for lags below ~200 ms. This large correlation coefficient may be explained by the effect of residual CI artifact leaking into the EEG. After 200 ms, the correlation coefficient drops rapidly and the curve presents two peaks. The morphology of the curves is consistent with previous studies in NH and CI users (Nogueira et al., 2019), and confirms that at later lags the effect of the artifact is reduced. Based on our previous work (Nogueira et al., 2019a; Nogueira et al., 2019b), we focus the analysis of the correlation coefficients on the lag interval ranging from 226 to 300 ms to minimize the effects of the CI artifact.

To investigate the influence of the training material, the statistical analysis was performed on the mentioned lag interval for the three different decoders (Decoder_HSM_, Decoder_Story_, Decoder_HSM+Story_). The rm-ANOVA revealed a significant effect of training material (F (2, 178) = 100.430, p < 0.0001) on the correlation coefficient to the attended speech stream. The results in Figure 1 show similar correlation coefficient values using either the Decoder_HSM_ or the Decoder_HSM+Story_. In contrast, the value of the correlation coefficient using the Decoder_Story_ was lower. For further analysis the Decoder_HSM_ was selected, because this decoder uses the same training material as the one used for testing the decoders. Moreover, this decoder provided the highest numerical correlation coefficients.

In Figure 2, it can be observed that at 0 dB SIR, the correlation coefficient to the attended speech stream was higher than the correlation coefficient to the unattended speech stream only for the Decoder_HSM_ and for the Decoder_HSM+Story_. AT this SIR condition, in theory, the CI artifact may have less impact on the correlation coefficients as the target and interference speech streams were delivered at the same presentation level. However, it cannot be discarded that one of the two voices (male or female) caused increased artifact explaining the larger correlation coefficient to the attended than to the unattended signal. This effect is investigated in section 4.2.2.

#### 4.2.2. EFFECT OF VOICE ON SELECTIVE ATTENTION DECODING

In this section we focus the analysis of the correlation coefficients using the Decoder_HSM_. During the experiment 20 lists were uttered by a male voice and 20 by a female voice. In order to assess if there was an effect of voice on the correlation coefficients, two additional decoders were designed. One decoder was trained using the female HSM sentences (Decoder_HSM_female_) and another one was trained using the male HSM sentences (Decoder_HSM_male_). The resulting correlation coefficients across lags for each decoder and SIR are presented in Figure 3.

**Figure 3:**
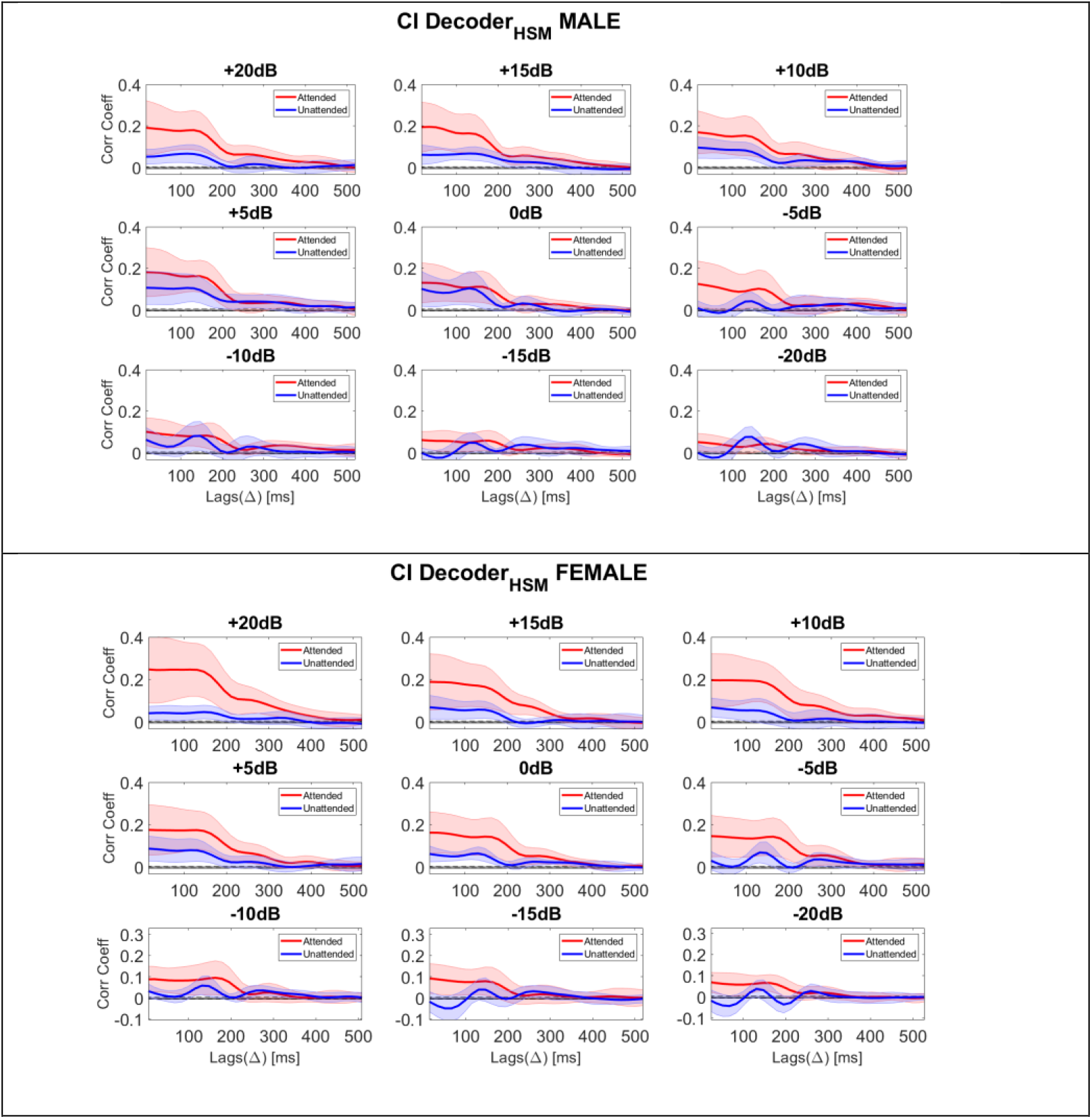
Correlation coefficients (*ρ_A_,ρ_U_*) to the attended and the unattended speech stream for cochlear implant (CI) users at different SIRs. The testing material used to compute the correlation coefficients consisted of the HSM sentence test uttered by a male (left) or a female (right) voice. Selective attention was decoded for SIRs ranging from −20 dB to +20 dB. The training material was based on the HSM sentence test to create the decoder Decoder_HSM_.

A 2-way-rm-ANOVA was conducted to assess if there were any significant effects of voice (male or female) and SIR on the averaged attended correlation coefficients in the lag range from 226 ms to 300 ms. The ANOVA showed no significant effect of gender (F(1, 9)= 4.375, p=0.058), a significant effect of SIR (F(8, 72)= 14.113, p<0.0001) and no significance interaction between both factors (F(8, 72)= 1.531, p=0.162). In general, the morphology of the correlation coefficient across lags does not present a substantial difference between the male and the female voice. Based on the outcomes of this statistical analysis, the correlation coefficients for both, the male and the female speech stream, were combined and the statistical difference between the attended and unattended correlation coefficient was investigated.

Figure 4 presents the correlation coefficients to the attended (*ρ_A_*) and to the unattended (*ρ_U_*) speech stream for CI users at different SIRs averaged across the lag interval 226-300 ms. An rm-ANOVA revealed a significant effect of attended vs unattended correlation coefficients (F (1, 9) = 13.885; p=0.005), a significant effect of SIR (F (8, 72) = 4.915; p = p<0.0001) and significant interaction effect of both factors (F (8, 72) = 9.957; p<0.0001). A post-hoc Wilcoxon signed-rank test with type 1 error Bonferroni correction revealed that for positive SIRs and for the SIR of −5 dB the correlation coefficient to the attended speech stream was larger than the correlation coefficient to the unattended speech stream (Figure 4). For the 0 dB condition, it can be observed that the correlation coefficient to the attended speech stream is larger than the correlation coefficient to the unattended speech stream, however the difference just missed significance after Bonferroni correction (p = 0.059).

**Figure 4:**
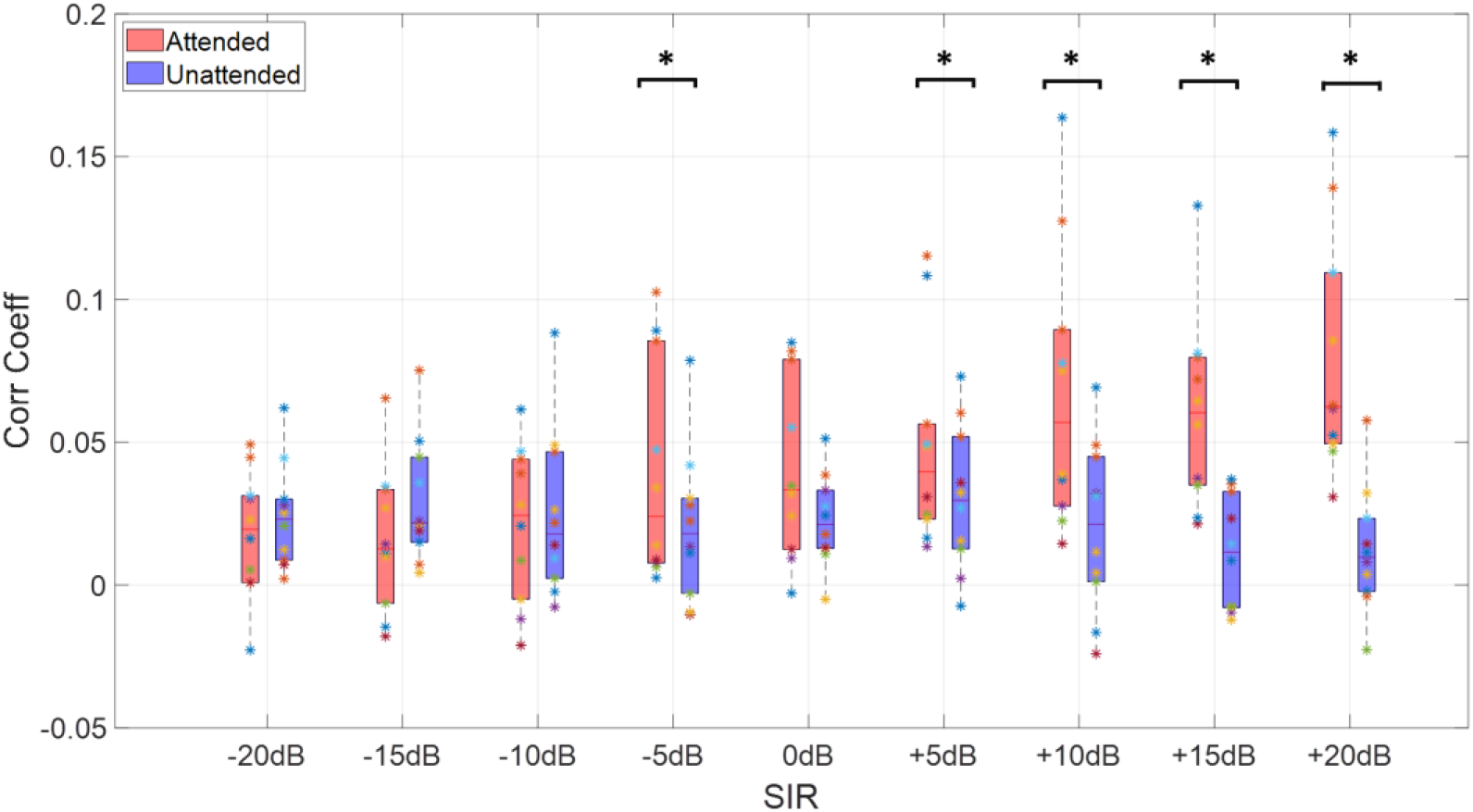
Correlation coefficients (*ρ_A_,ρ_U_*) to the attended and to the unattended speech stream for cochlear implant (CI) users at different signal to interference ratios (SIRs) averaged across the lag interval 226-300 ms. The asterisks denote the pairs (attended vs unattended correlation coefficients) that were significantly different for each SIRs. The statistical test was conducted using post-hoc pairs Wilcoxon test.

It is important to remark that if the artifact would be the only factor contributing to the attended correlation coefficient, one would expect the unattended correlation coefficient to be higher than that the attended correlation coefficient at negative SIRs. However, this was not the case and for SIR=-5 dB the attended correlation coefficient was significantly higher than the unattended correlation coefficient as revealed by a Wilcoxon signed ranked test with Bonferroni type I error correction (p = 0.037). From these results, it can be concluded that it is possible to decode selective attention when both speech streams are presented monaurally to CI users.

#### 4.2.3. EFFECT OF SIGNAL TO INTERFERENCE RATIO (SIR) ON SELECTIVE ATTENTION DECODING

This section investigates the attended *ρ_A_SIR__* and unattended *ρ_U_SIR__* correlation coefficients across SIRs as well as the difference between both correlation coefficients (*ρ_A_SIR__ – ρ_U_SIR__*). Note that the SIR ranged from −20 to +20 dB and at positive SIRs the level of the target signal was kept constant while the level of the interference was reduced. In contrast, for negative SIRs, the level of the interferer was kept constant while the level of the target was decreased. Hence, the correlation coefficients *ρ_A_SIR__* and *ρ_U_−SIR__* were computed when the attended and the unattended signal were presented at the same level. the If residual CI artifact leaks into the EEG, it is expected that the correlation coefficient to the attended speech or to the difference between both correlation coefficients will increase with SIR. For this reason, we also analyzed the difference between the attended correlation coefficient at a given SIR and the unattended correlation coefficient at the opposite SIR (*ρ_A_SIR__* – *ρ_U_−SIR__*). We hypothesized that difference in correlation coefficients at opposite SIRs balances the effect of residual CI electrical artifact at different SIRs.

The correlation coefficient to the attended speech stream (*ρ_A_SIR__*; top left panel) and to the unattended speech stream (*ρ_U_SIR__*; top right panel), as well as the difference correlation coefficient between the attended and unattended speech stream at the same (*ρ_A_SIR__* – *ρ_U_SIR__*; bottom left panel) and at opposite SIRs (*ρ_A_SIR__* – *ρ_U_−SIR__*; bottom right panel) averaged across lags ranging from 226-300 ms are presented in Figure 4. The grey shaded areas in Figure 5 denote the chance correlation coefficient level which was estimated from the correlation coefficients between the reconstructed speech stream for a particular HSM list and the envelope speech stream of other randomly selected HSM lists. The chance correlation coefficient level was estimated from 40 HSM lists (20 uttered by the male and 20 uttered by the female voice) corresponding to 40 folds of the cross-validation procedure.

**Figure 5:**
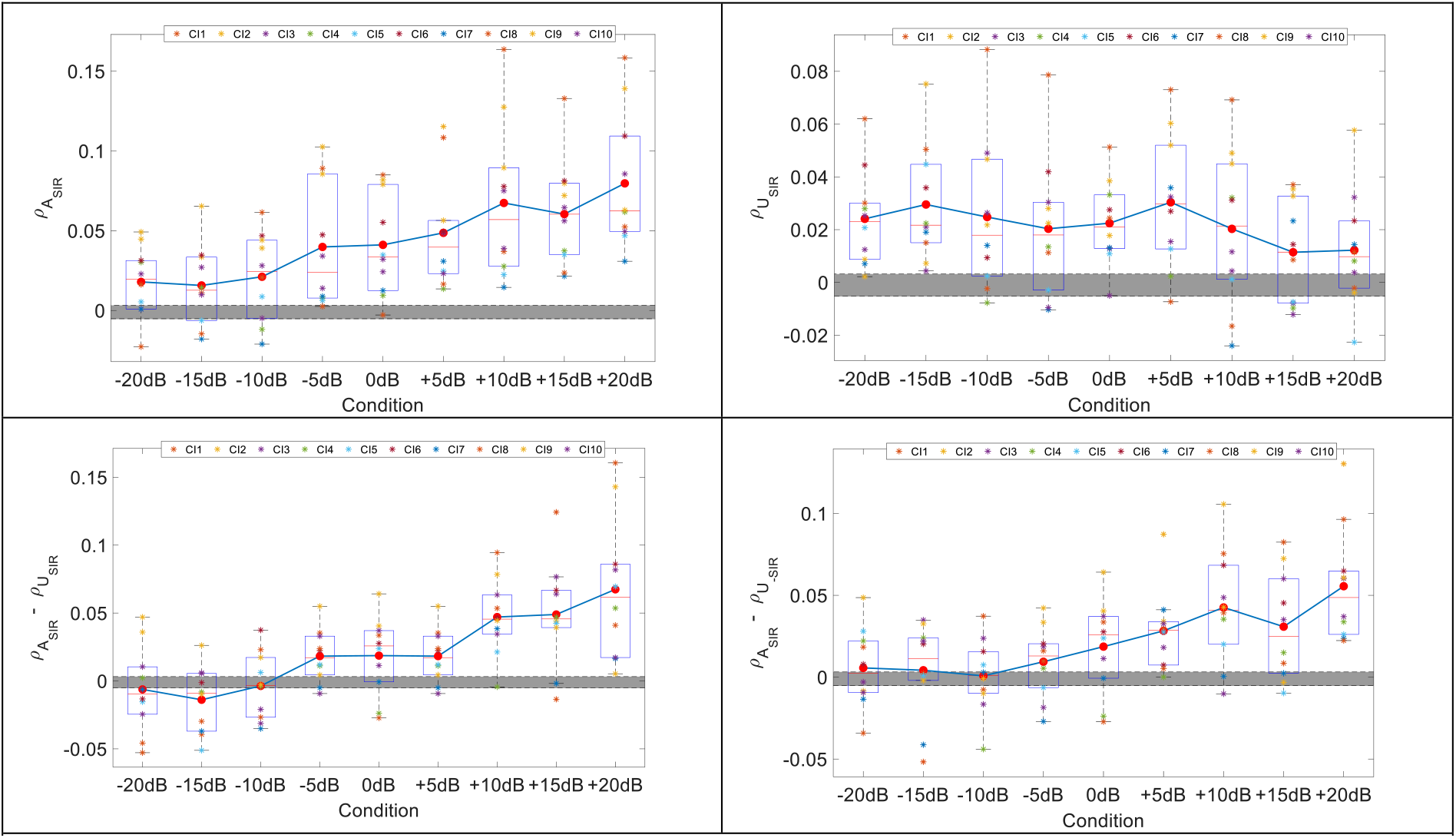
Correlation coefficients averaged across lags ranging from 226 ms to 300 ms at different signal to interference ratios (SIRs) measured in cochlear implant (CI) users for the attended speech (*ρ_A_SIR__*; top left); for the unattended speech (*ρ_U_SIR__* top right); for the difference between the attended and unattended correlation coefficients at the same (*ρ_A_SIR__ – ρ_U_SIR__*; bottom left) and opposite SIRs (*ρ_A_SIR__* – *ρ_U_−SIR__*; bottom right). The grey shaded area denotes the chance correlation coefficient level.

The results in Figure 5 show that *ρ_A_SIR__*, *ρ_A_SIR__* – *ρ_U_SIR__*, and *ρ_A_SIR__* – *ρ_U_−SIR__* become larger with increasing SIR. A one way repeated measures of ANOVA confirmed a significant effect of SIR on the *ρ_A_SIR__* (F (8, 72) = 14.113; p<0.0001), on *ρ_A_SIR__* – *ρ_U_SIR__*(F (8, 72) = 10.559; p<0.0001) and on the *ρ_A_SIR__* – *ρ_U_−SIR__* (F (8, 72) = 6.546; p=0.0001), but not on the unattended correlation coefficient *ρ_U_SIR__* (F (8, 72) = 1.165; p=0.342). It is important to remark that the difference correlation coefficient at opposite SIR (*ρ_A_SIR__* – *ρ_U_−SIR__*), is the least influenced by any CI artifact and it significantly increases with SIR.

Following the ANOVA, a correlation analysis between the correlation coefficients and the SIR was computed. *ρ_A_SIR__, ρ_A_SIR__ – ρ_U_SIR__*, and *ρ_A_SIR__ – ρ_U_−SIR__* significantly correlated with SIR with correlation coefficients and p values (r = 0.5086, p=0.001, (r = 0.3702, p = 0.0003) and (r = 0.3585, p = 0.0005), respectively. No significant correlation between *ρ_U_SIR__* and SIR was observed (r = 0.037; p = 0.7241).

### 4.3. CORRELATION BETWEEN SELECTIVE ATTENTION AND SPEECH UNDERSTADING PERFORMANCE IN COCHLEAR IMPLANT USERS

Figure 6 presents the correlation analysis between speech understanding scores in percentage of correct words and the selective attention correlation coefficients: *ρ_A_SIR__,ρ_U_SIR__, ρ_A_SIR__ – ρ_U_SIR__*, and *ρ_A_SIR__ – ρ_U_−SIR__* for each individual subject across SIRs. The speech understanding performance was measured only at positive SIR conditions, as the negative SIR conditions were too challenging for CI users and the speech score would have been zero percent.

**Figure 6:**
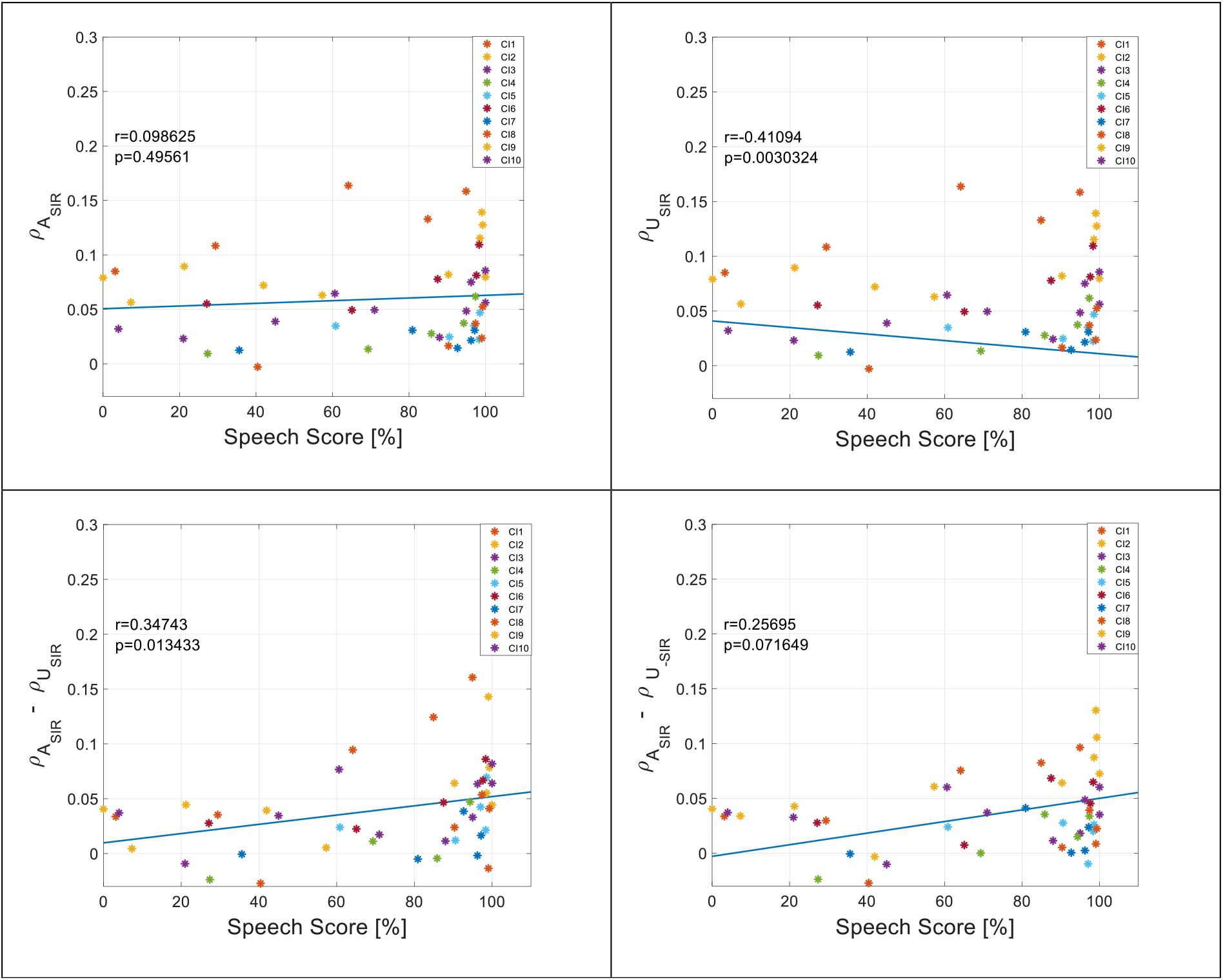
Correlation between the speech understanding scores using the HSM sentence test and the correlation coefficient to the attended speech (*ρ_A_SIR__*; Top left panel), to the unattended speech (*ρ_A_SIR__*; Top right panel) and to the difference correlation coefficient between the attended and the unattended speech stream at the same SIR (*ρ_A_SIR__ – ρ_U_SIR__*; Bottom left panel) and at opposite SIRs (*ρ_A_SIR__ – ρ_U_−SIR__*; Bottom right panel).

The results from Figure 6 show that in general the correlation coefficients tend to become higher with increasing speech understanding performance, except for *ρ_U_SIR__* which shows the opposite trend. However, the only significant correlation between speech understanding scores and selective attention decoding was observed for the correlation coefficients *ρ_A_SIR__ – ρ_U_SIR__* (r = 0.34; p = 0.013) and for *ρ_U_SIR__* (r = 0.41; p = 0.003) after correcting for type I error for multiple comparisons (N=3 resulting in p<0.016).

## 5. DISCUSSION

This work investigated the possibility to decode selective attention in CI users when two speech streams are presented monaurally and at different SIRs. Moreover, this study investigated the influence of using different types of training and test material on the correlation coefficients derived from a selective attention paradigm. The end goal of the present work was to investigate whether behavioral speech understanding performance can be predicted from the selective attention correlation coefficients at different SIR levels.

### 5.1 Selective attention decoding to monaural presentation of speech in cochlear implant users

Previous research showed that it is possible to decode selective attention in NH listeners through dichotic (e.g. O’Sullivan et al., 2012; Mirkovic et al., 2015) and monaural presentation of two concurrent speech streams (e.g. Ciccarelli et al., 2019; Mesgarani et al., 2012). In CI users, Sommers et al. (2019) showed that it is possible to conduct EEG neural tracking when a single monaural speech stream with background noise is used. Moreover, previous studies showed the possibility to decode selective attention in bilateral CI users through dichotic presentation of two concurrent speech streams (Nogueira et al., 2019a; 2019b) or by presenting each speech stream through an independent loudspeaker (Paul et al., 2020). The current study extends these previous works and shows that it is also possible to decode selective attention when both speech streams are monaurally presented to CI users. Moreover, selective attention was decoded at positive and negative SIRs computing the correlation coefficients between the reconstructed speech and the attended or the unattended speech stream. The results of the current study show that the attended correlation coefficient presented two peaks at around 70 and 180 ms, for CI users (Figure 2) and for NH listeners (Figure 8 of Appendix 2). These peaks became smoother with decreasing SIR. Results from our previous studies (Nogueira et al., 2019a; 2019b) show that these two peaks were only observed for NH listeners and not for CI users when the SIR was 0 dB. Therefore, the appearance of these two peaks may be caused by the use of higher SIR conditions in the current study which made the task easier for the CI users. In agreement with the same previous works (Nogueira et al., 2019a; 2019b), high correlation coefficients in the range of 0.1-0.4 were observed at early lags. The correlation coefficients at these early lag intervals are probably more contaminated by the electrical artifact. The use of later lags was also suggested by the TRF analysis performed by Paul et al. (2020), who showed that late cortical responses (around 250 ms) are more suitable to decode selective attention in comparison to early cortical responses at 150 ms in bilateral CI users. In order to reduce the effect of the artifact, ICA artifact rejection, as suggested by Viola et al. (2012), was used. Moreover, we focused our analysis on the correlation coefficients at later lags (226-300 ms) minimizing the effect of the CI artifact (Nogueira et al., 2019a; 2019b).

The results of the current study showed significantly higher correlation coefficients to the attended speech stream than to the unattended speech stream in CI users, especially for positive SIRs. At positive SIRs, the level of the attended speech stream was kept fixed while the level of the unattended speech stream was decreased. For this reason, it was expected that with higher SIR, the correlation coefficient to the attended speech stream would increase as the person may be able to focus better and the correlation coefficient to the unattended speech stream would decrease. The reason being that residual CI electric stimulation leaks into the EEG, even if ICA artifact removal was applied and late lags were used for the analysis. However, the results show that the attended correlation coefficient increased significantly with SIR (p<0.0001). In contrast, for negative SIRs, the presentation level of the unattended speech stream was fixed and the level of the attended speech stream was decreased. For this reason, it is remarkable that for SIR=-5dB, the correlation coefficient to the attended speech stream was significantly larger than to the unattended speech stream, for both the male and the female decoder. These results demonstrate that it is possible to decode selective attention from EEG with monaural presentation of two concurrent speech streams in CI users even at a negative SIR.

### 5.3 Factors influencing selective attention decoding

#### Effect of Training Material

In previous studies, the speech materials used for training and testing the decoder were based on the same story books (e.g. O’Sullivan et al., 2012; Mirkovic et al., 2015; Nogueira et al., 2019a; 2019b) or the same digit-in-noise test (Paul et al., 2020). In contrast, Lesenfants et al. (2019) used a story book uttered by a single voice for training the decoder and sentences in noise to test the decoder. In the current work, the effect of using different speech materials for training and testing the decoders was investigated, three decoders were designed for this purpose: the first one was trained using a story book (Decoder_Story_), the second one was trained based on HSM sentences (Decoder_HSM_) and the third one was trained based on both, the story book and the HSM sentences (Decoder_HSM+Story_). The three decoders were tested using HSM sentences at different SIRs. The results of the investigation show that the training material had a significant effect on the attended correlation coefficient (p < 0.0001). The Decoder_Story_ used the most different materials between training and testing and resulted in the lowest attended correlation coefficients. It is interesting to note that with the Decoder_Story_, the attended correlation coefficient gradually became higher with increasing SIR (Figure 2), probably because the training material used to create this decoder was more similar to the test material conditions with higher SIRs. To avoid this bias in the decoders, we continued the selective attention decoding analysis based on the Decoder_HSM_ which uses the same type of speech material and with balanced SIRs for training and testing the decoder.

#### Effect of voice

The selective attention paradigm used in the current study was based on two concurrent speech streams, one uttered by a male voice and one by a female voice. Two DecodersHSM were created, one based on the male voice and one based on the female voice. The analysis of the results showed no significant difference between both decoders. These results indicate that selective attention can be decoded in CI users irrespective of the speech stream attended by the listener. For this reason, the following results were analyzed creating decoders that used both voices.

#### Effect of Signal to Interference Ratio

It is know<u>n</u> that the CI introduces and electrical artifact that leaks into the EEG and that this artifact is related to the speech envelope. However, the artifact produced by speech signals is weaker than the artifact introduced by more deterministic stimuli such as pulse trains or pure tones (Wagner et al., 2018). An advantage of the selective attention paradigm w.r.t. neutral tracking paradigms to single stimuli is that it can be used to analyze the differences between the attended and the unattended correlation coefficients. Assuming similar artifact contributions from both speech streams and after using late lags for the analysis as well as ICA artifact removal, it is expected that the differences between both correlation coefficients are dominated by neural activity rather than by the CI artifact. The effect of SIR on selective attention decoding was investigated in previous research (Lesenfants et al., 2019) and showed higher correlation coefficients with increasing SIR in NH listeners. In the present work, the attended correlation coefficient measured in CI users became higher with increasing SIR (p<0.0001), which confirms this previous work. Moreover, the attended and the unattended correlation coefficient difference became larger with increasing SIRs (p<0.0001). However, it is possible that this increase in the attended correlation coefficient (*ρ_A_SIR__*) or the difference between the attended and the unattended correlation (*ρ_A_SIR__ – ρ_U_−SIR__*) coefficients is caused by residual CI leaking into the EEG. The reason being that positive SIRs were created by reducing the level of the unattended speech stream while keeping constant the level of the attended speech stream, and hence the contribution of the unattended speech to the CI artifact will be less with increasing SIR. For this reason, we introduced a new method that may be less influenced by the CI artifact. The new method uses the difference between the attended and the unattended correlation coefficients at opposite SIRs (*ρ_A_SIR__ – ρ_U_−SIR__*). Negative SIRs were created keeping the level of the unattended speech and reducing the level of the attended speech. Hence the subtraction of the attended and unattended correlation coefficients at opposite SIRs should balance any residual effect of the CI artifact. The results of the current study show that this difference between correlation coefficients increased with greater SIR (p=0.0001).

### 5.6 Relation between Speech Understanding and Selective Attention Decoding

A goal of the present study was to predict speech understanding performance based on the correlation coefficients derived from the selective attention decoding paradigm. Paul et al. (2020) showed that it is possible to predict speech understanding performance of CI users from the TRF at a fixed SIR when late cortical responses (250 ms) are used, but these results could not be confirmed in NH listeners. The present study measured behavioral speech understanding performance and selective attention decoding presenting two concurrent speech streams at different SIRs using the same speech material for both tasks. The speech performance of CI users was measured only at positive SIRs, as the negative SIR conditions were too challenging for them. The results of the present study show no significant correlation between the attended correlation coefficient and behavioral speech understanding performance. However, a significant correlation was observed between the behaviorally measured speech understanding performance and the difference between the attended and the unattended correlation coefficients (r = 0.34; p = 0.013). However, it cannot be completely excluded that this result is explained by residual CI artifact, even if ICA rejection was applied and late lags were used for the analysis. This hypothesis is confirmed by the significant negative correlation between the unattended correlation coefficient and the behaviorally measured speech understanding performance (r = −0.41; p = 0.003). In contrast, the difference between the attended and the unattended correlation coefficients at opposite SIRs (*ρ_A_SIR__ – ρ_U_SIR__*) which in principle balances the contribution of the CI artifact did not significantly correlate with speech understanding performance, although a small trend was observed (r = 0.25; p = 0.07).

## 6. CONCLUSION

The results of the study confirm that it is possible to decode selective attention in CI users even if continuous artifact is present in the recordings. Moreover, the results show that it is possible to decode selective attention when two concurrent speech streams are presented monaurally to CI users. The study demonstrates that the selection of the type of material for training the selective attention decoder significantly influences the results. In general, it was shown that the correlation coefficient to the attended speech stream or the difference between the attended and the unattended correlation coefficients increase with SIR. However, it cannot be completely excluded that these effects are caused by residual CI artifact leaking into the EEG. The current study presents a novel measure based on the difference between the attended and the unattended correlation coefficients at opposite SIRs which further balances the residual CI artifact. However, this measure only showed a correlation trend with the speech understanding performance of the CI users.

## APPENDIX 1: DECODING SELECTIVE ATTENTION AT DIFFERENT SIGNAL TO INTERFERENCE RATIOS WITHOUT ARTIFACT REJECTION

Figure 6 presents the attended and the unattended correlation coefficients of selective attention before applying Infomax ICA artifact rejection averaged across the CI users that participated in the study. Note that positive SIRs were obtained by setting the level of the target speaker and reducing the level of the interferer. In contrast, negative SIRs were obtained by setting the level of the interferer and reducing the level of the target speaker. In Figure 6, it can be seen that for negative SIRs, the unattended correlation coefficients are higher than the attended correlation coefficients. This effect can be explained by the higher level of the target stream.

The use of ICA significantly decreases the unattended correlation coefficients, but not the attended correlation coefficients at the positive SIRs. It is also to observe that at 0 dB SIR, where both streams were presented at the same level, the attended correlation coefficients are higher than unattended.

Figure 8 presents the attended and unattended correlation coefficients at different SIR for CI users. Note, that here the data without ICA rejection is presented. The post-hoc Wilcoxon Test was conducted and revealed significance for all SIR pairs of attended and unattended correlation coefficients except SIRs of −10 dB (p = 0.059) and −5 dB (p = 0.241). Comparing to Figure 3, ICA rejection manifested the suppression of unattended speech stream at negative conditions, while attended at positive condition stayed almost without changes. The effect of suppression showed especially the SIR of −5dB, as after ICA rejection the difference between attended and unattended became significant (p = 0.22). It was observed that before ICA rejection the SIR of 0 dB showed the significance (p = 0.022) while after rejection this difference was insignificant (p=0.59). As this SIR is less contaminated by the artifact, because speech streams level was maintained the same for attended and unattended, this cannot be explained by the artifact.

**Figure 7:**
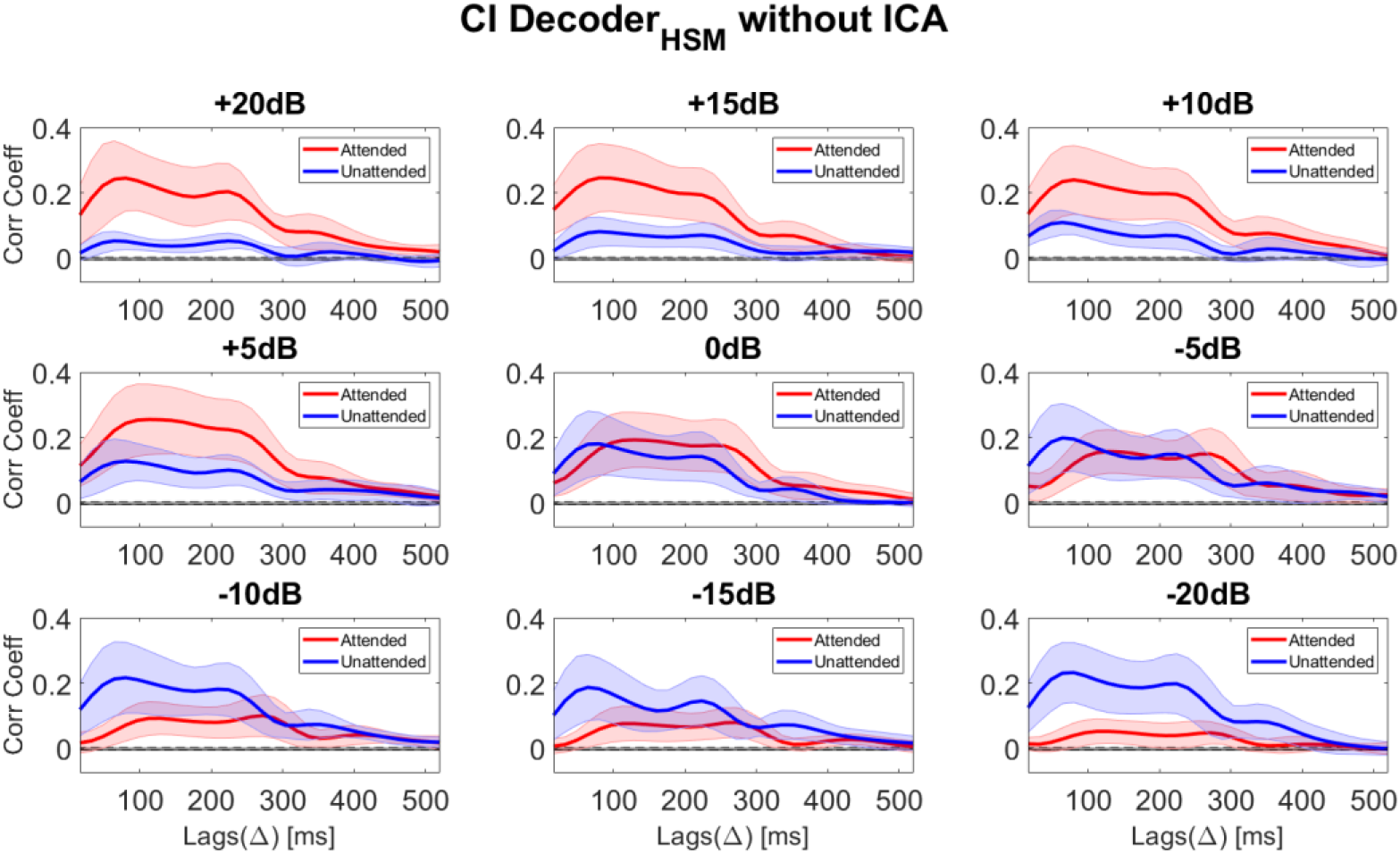
Correlation coefficients (Corr Coeff) to the attended and the unattended speech stream without ICA artifact rejection for Cochlear implant (CI) users at different SIRs. Selective attention was decoded for SIRs ranging from −20 dB to +20 dB SIR.

**Figure 8:**
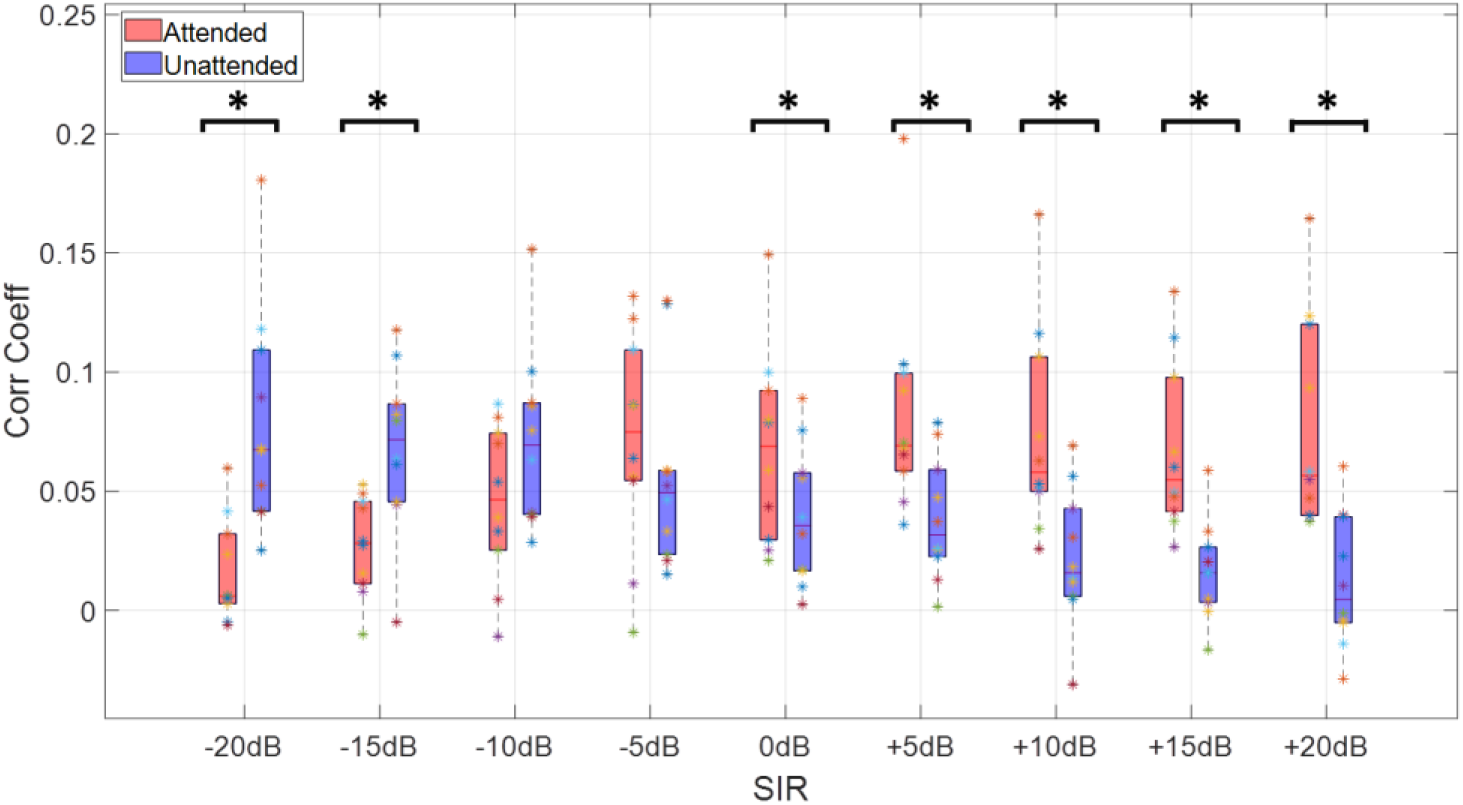
Correlation coefficients (*ρ_A_,ρ_U_*) to the attended and the unattended speech stream for cochlear implant (CI) users at different SIRs at the lag interval 226-300 ms before applying ICA rejection method. With asterisks marked the pairs of SIRs which showed significant difference between attended and unattended correlation coefficients. The statistical test was conducted using post-hoc pairs Wilcoxon test.

## APPENDIX 2: DECODING SELECTIVE ATTENTION IN NORMAL HEARING LISTENERS

The same measures performed in CI users were also conducted in a control group of 5 NH listeners (mean age = 26.8). All NH participants did not have hearing loss. The EEG recordings were obtained using exactly the same methodology as in CI users but presenting the two speech streams via an insert earphone (3M E-A-RTONE 3A, 50 Ohm) monaurally. Each participant adjusted the presentation level to a moderate loudness level by means of a seven-point loudness-rating scale (where 1 is “very soft” and 7 is “extremely loud”). The attended and the unattended correlation coefficients were calculated for the Decoder_HSM_ at 9 different SIRs (Figure 8).

In Figure 9, it can be observed that at early lags, the correlation coefficients for NH listeners presented lower values than for CI users (around 0.04 for NH listeners compared to 0.25 for CI users; Figure 7). This large difference may be explained by the contribution of the CI artifact at early lags. Moreover, previous works show larger correlation coefficients for hearing impaired subjects compared to NH listeners (Decruy et al., 2020). The morphology of the curves is consistent with previous works of Nogueira et al. (2019a; 2019b) showing two distinguished peaks at lags corresponding with 120 ms and 250 ms. The second peak in the attended correlation coefficient curve can also be observed in the curves obtained in CI users. For this reason, the same lag interval ranging from 225 ms to 300 ms was used in the analysis.

**Figure 9:**
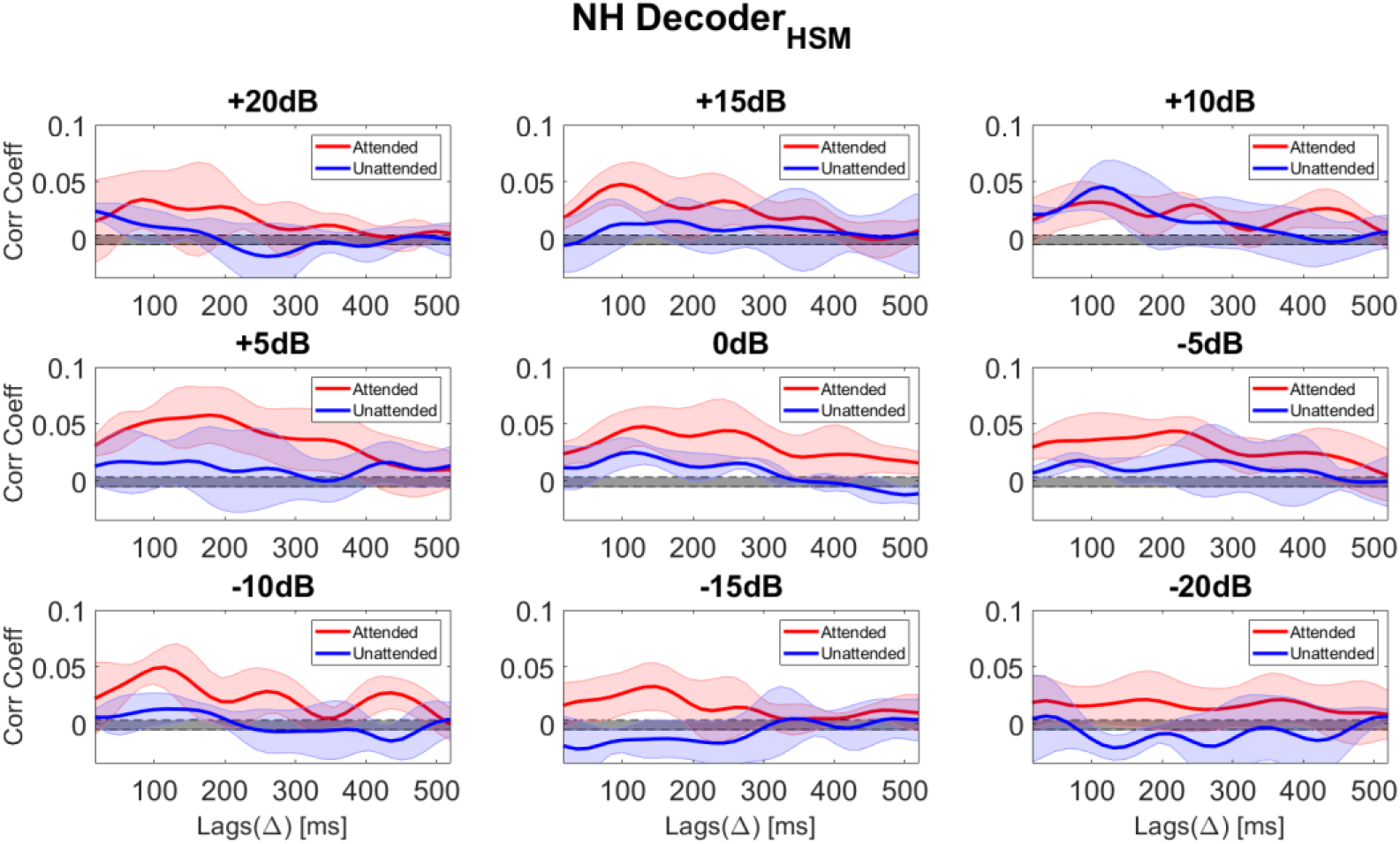
Attended and unattended correlation coefficients (Corr Coeff) without ICA artifact rejection for normal hearing listeners (NH) at different SIRs ranging from −20 dB to +20 dB.

## DATA AVAILABILITY

All data can be made available upon request to the authors.

## ETHICAL STATEMENT

The study was carried out in accordance with the declaration of Helsinki principles and approved by the ethics committee of the Hannover Medical School (Hanover, Germany). The participants provided their written informed consent to participate in this study.

## CONFLICT OF INTEREST STATEMENT

The authors declare that the research was conducted in the absence of any commercial or financial relationships that could be construed as a potential conflict of interest.

## ACKNOWLEDGEMENTS

We would like to thank all participants for their time and patience. This work was supported by the Deutsche Forschungsgemeinschaft (DFG, German Research Foundation) under Germany’s Excellence Strategy EXC 2177—Project ID 390895286.

## Footnotes

1 In Appendix 1 the same data is presented before applying ICA artifact rejection.

## Notes

### Competing Interest Statement

The authors have declared no competing interest.

